# DiffBrainNet: differential analyses add new insights into the response to glucocorticoids at the level of genes, networks and brain regions

**DOI:** 10.1101/2022.04.21.489034

**Authors:** Nathalie Gerstner, Anthi C. Krontira, Cristiana Cruceanu, Simone Roeh, Benno Pütz, Susann Sauer, Monika Rex-Haffner, Mathias V. Schmidt, Elisabeth B. Binder, Janine Knauer-Arloth

## Abstract

Genome-wide gene expression analyses are invaluable tools for increasing our knowledge of biological and disease processes, allowing a hypothesis-free comparison of gene expression profiles across experimental groups, tissues and cell types. Traditionally, transcriptomic data analysis has focused on gene-level effects found by differential expression. In recent years, network analysis has emerged as an important additional level of investigation, providing information on molecular connectivity, especially for diseases associated with a large number of linked effects of smaller magnitude, like neuropsychiatric disorders and their risk factors, including stress. In this manuscript, we describe how combined differential expression and prior-knowledge-based differential network analysis can be used to explore complex datasets. As an example, we analyze the transcriptional responses following administration of the glucocorticoid/stress hormone receptor agonist dexamethasone in *C57Bl/6* mice, in 8 brain regions important for stress processing: the prefrontal cortex, the amygdala, the paraventricular nucleus of the hypothalamus, the cerebellar cortex, and sub regions of the hippocampus: the dorsal and ventral *Cornu Ammonis* 1, the dorsal and ventral dentate gyrus. By applying a combination of differential network- and differential expression-analyses, we find that these explain distinct but complementary aspects and biological mechanisms of the responses to the stimulus. In addition, network analysis identifies new differentially connected partners of important genes and can be used to generate hypotheses on specific molecular pathways affected. With this work, we provide an analysis framework and a publicly available resource for the study of the transcriptional landscape of the mouse brain: DiffBrainNet (http://diffbrainnet.psych.mpg.de), which can identify molecular pathways important for basic functioning and response to glucocorticoids in a brain-region specific manner.

## Introduction

High-throughput transcriptomics are extensively employed to study healthy as well as disease-related tissue expression profiles from *in vitro* and *in vivo* model systems or human tissue. Traditionally, transcriptomic data analysis has been based on differential expression (DE) analysis and has focused on gene-level associations to phenotypes. In the last decade, gene set enrichment analysis [1] and network analysis [2–4] have emerged allowing the study of complex associations between sets of genes, in multiple tissues and for multiple outcomes [5,6,15–17,7–14].

Network analysis is critical for the study of relationships between genes, and in turn, of molecular pathways. This is especially true for complex disorders for which risk is conferred by a combination of many small effects. Strong DE can be expected with major genetic or environmental impacts such as in cancer [18,19]. For other disorders, for example in neuropsychiatry, risk is driven by multiple polygenic and interlaced environmental factors that affect a multitude of transcripts, often with only small effect sizes [20,21]. A combinatorial analysis framework of DE and network analysis has proven very useful for unraveling additional biology and pathomechanisms of complex disorders [5]. For example, gene co-expression networks, based on Pearson correlations, along with DE analysis have been used to study shared and distinct transcriptomic profiles of five major neuropsychiatric disorders (autism spectrum disorder; schizophrenia, bipolar disorder, major depressive disorder and alcoholism) leading to the identification of gene modules associated with specific cell-types and disorders [22].

Besides correlation-based methods, which tend to suffer from over-connectivity and low specificity, several other classes of algorithms are used for network inference [23]. More advanced are, for example, regression-based or Bayesian methods. While Bayesian methods perform poorly on large datasets and are more suitable for small networks [23], regression- and other machine learning-based algorithms require large amounts of samples to confidently infer connections in a high-dimensional input space. To overcome this limitation of regression-based network inference and increase the performance on datasets with small amounts of samples, the input space can be reduced by facilitating prior-knowledge [24]. Prior-knowledge refers to already described functional relationships between transcripts or proteins, accessible from publicly available databases. The Knowledge guided Multi-Omics Network inference approach (KiMONo) implements such a combination of prior-guided regression-based network inference and was previously shown to be a powerful approach to infer integrated multi-level networks [3].

Traditionally the stimulus or disease impact on networks has been modeled by associating modules of co-expressed genes with disease phenotypes or comparing the number of connections a gene has in the control and stimulus networks. This has proven challenging given that it is based on the comparison of networks with different topological characteristics [4]. To tackle this, differential network (DN) analysis has emerged. DN analysis computes the differential co-expression and regulatory interactions of many genes in a single network and analyzes biological processes inferred from one DN [25], thus eliminating the problems arising when trying to compare two or more networks at different stimulation paradigms. DN analysis offers the possibility to study the directed multivariate effects of the treatment or disease state on the genes’ neighborhoods. Another advantage of using prior-knowledge network analysis algorithms is that the inferred networks have the same topological characteristics which results in a more robust calculation of the differential connections.

In this study, we now leverage the power of DN approaches and calculate regression- and prior-knowledge-based genome-wide networks from RNA expression data of 8 mouse brain regions following a vehicle or a pharmacological stimulus, and compute differential networks in addition to differential expression. As a stimulus we used dexamethasone, a synthetic glucocorticoid that is a preferential agonist of the glucocorticoid receptor (GR). GR is a transcription factor able to elicit a robust transcriptomic response when bound to its agonists [26], it is an important component of the stress-axis and has been implicated in risk for stress-related psychiatric disorders [27]. The 8 brain regions were selected for their implication with the activation of the stress axis and the response to stress, and include a detailed segmentation of the hippocampal formation (ventral and dorsal dissections of both *Cornu Ammonis* 1-CA1 and dentate gyrus-DG), the prefrontal cortex (PFC), the amygdala (AMY), the cerebellar cortex (CER) and the paraventricular nucleus of the hypothalamus (PVN). We combined DN with DE analysis in order to provide an analysis framework for transcriptomic data and a resource of all levels of information. This public resource is named DiffBrainNet (DiffBrainNet access: http://diffbrainnet.psych.mpg.de). We provide examples of how DiffBrainNet can be used to study the molecular landscape of the brain and unravel biological mechanisms of response to dexamethasone and GR activation in a brain region-specific manner.

## Materials and Methods

### Experimental animals

All experiments and protocols were performed in accordance with the European Communities’ Council Directive 2010/63/EU and were approved by the committee for the Care and Use of Laboratory animals of the Government of Upper Bavaria. All mice were obtained from the in-house breeding facility of the Max Planck Institute of Psychiatry and kept in group housed conditions in individually ventilated cages (IVC; 30cm × 16 cm × 16 cm; 501 cm2) serviced by a central airflow system (Tecniplast, IVC Green Line – GM500). Animals had ad libitum access to water (tap water) and standard chow and were maintained under constant environmental conditions (12:12 hr light/dark cycle, 23 ± 2 °C and humidity of 55%). All IVCs had sufficient bedding and nesting material as well as a wooden tunnel for environmental enrichment. Animals were allocated to experimental groups in a semi-randomized fashion, data analysis and execution of experiments were performed blinded to group allocation. 3-months old C57Bl/6n male mice (n=15 animals per condition) were injected intraperitoneally with dexamethasone at a dose of 10 mg/kg body weight (treatment) or 0.9% saline as control (vehicle). Four hours later the mice were sacrificed, the brain was perfused with a solution of Heparin in 0.9% saline, extracted and snap-frozen in butanol on dry ice and kept in -80°C until further use. The brains were cut in 250μm coronal slices and 8 brain regions were isolated following the stereotaxic coordinates of the mouse brain atlas [28]. In detail, the following brain regions were isolated: cingulate cortex 1 and 2 (bregma 2.34 to -0.22), from now on referred-to as prefrontal cortex (PFC); paraventricular nucleus of the hypothalamus (PVN; bregma -0.58 to -1.22); amygdala (AMY; bregma 0.02 to -0.94); dorsal Cornu Ammonis 1 (dCA1; bregma -1.22 to -2.80); ventral Cornu Ammonis 1 (vCA1; bregma -2.92 to -3.88); dorsal dentate gyrus (dDG; bregma -0.94 to -2.80), ventral dentate gyrus (vDG; bregma -2.92 to -3.88) and cerebellar cortex (CER; bregma -5.80 to -6.24). Brain punches were kept in dry ice while cutting and then in -80oC until the RNA extraction was performed.

### RNA extraction

RNA was extracted using an automated Chemagic 360° instrument with an integrated dispenser and the chemagic RNA Tissue Kit (CMG-1212) following manufacturer’s instructions. In short, Chemagic 360° RNA extraction is based on the use of magnetic beads that bind the nucleic acids which are then isolated using magnetized metal rods. Homogenization of the tissue was achieved using rotating zirconium beads. Washing steps and subsequent elution of the RNA was achieved by switching off the magnet while the rods continue to rotate in a buffer of preference. DNA was digested using DNase I and proteins using Proteinase K. RNA concentration was measured using a Nanodrop and the quality was measured using Tapestation RNA ScreenTapes (High Sensitivity RNA ScreenTapes, Cat No. 5067-5579).

### RNA sequencing

3’ tag RNA sequencing libraries were prepared using the QuantSeq 3’ mRNA Fwd kit (Lexogen) following manufacturer’s instructions with the addition of unique molecular identifiers (UMIs-UMI Second Strand Synthesis Module for QuantSeq FWD) for the tagging of individual transcripts. Libraries were single-end sequenced on an Illumina HiSeq4000 sequencer using 75bp long reads for a total coverage of an average of 10M reads per library. Five samples were excluded from sequencing and/ or further analysis due to technical issues with the library preparation: two dexamethasone-treated dCA1 samples, one dexamethasone-treated PFC sample, one control PVN sample and one control vCA1 sample.

### RNA sequencing analysis

The quality of sequencing data was analyzed with FastQC v0.11.4 [29] and adapter trimming was performed with cutadapt v1.11 [30]. Unique molecular identifiers were extracted with UMI-tools v.0.5.4 [31], before the reads were aligned with the mouse reference genome (mm10, Ensembl release 84) using STAR v2.6.0a [32]. Afterwards, reads were deduplicated with UMI-tools and gene expression was quantified with featureCounts v1.6.4 [33]. The subsequent analysis was performed in R version 4.0.5 [34]. All genes that were not detected in at least one full treatment group were removed from the dataset leaving 12,976 genes. Subsequently, genes with less than 10 counts across all samples within each brain region were excluded (detailed numbers of genes per brain region in Table S1). To identify outliers, we performed a principal component analysis (PCA) on the samples of each brain region and treatment group separately. Samples with a distance of more than 2.5 standard deviations from the mean in the first principle component were excluded (numbers of outliers per brain region and treatment group in Table S1). Surrogate variable analysis (SVA)[35] was applied to account for unwanted variation in the data.

### Differential expression (DE) analysis

Significant surrogate variables (exact numbers in Table S1) were included as covariates in the DE analysis. The expression data was normalized and transformed using the vst function of DESeq2 v1.30.1 [36] for SVA and subsequent network analysis. DE analysis between the two treatment groups was performed for each brain region individually. We tested for DE with DESeq2 using the Wald test and reported the genes with a false discovery rate (FDR) below 10% as significant.

### DiffBrainNet

#### Network inference

Networks were generated for vehicle-(referred-to as “vehicle”) and dexamethasone-treated (referred-to as “treatment”) samples separately for each brain region using the network inference method: KiMONo [3]. KiMONo uses prior information from existing biological databases that provide the edges among the transcripts, as a basic network layout. Different omic layers (here only transcriptomic data) are then used on top of the prior basic-network layout to fit the edge weights in the network. Edge weights can thereby take on a value smaller than a predefined threshold which leads to the removal of the edge from the network (Fig 1). More specifically, KiMONo uses a multivariate regression approach with sparse group lasso penalization to model the expression levels of the transcripts. The possible predictors in the regression model are inferred from the gene’s connections in a prior network. In the inferred directed gene expression networks, the nodes represent transcripts of the input data and the edge weights are the beta coefficients (β-value) fitted by the regression approach (S1B Fig). A β-value > 0 indicates that two genes’ expression levels are correlated positively, while a β-value < 0 indicates that two genes’ expression levels are correlated negatively. Significant surrogate variables identified during DE analysis were used as covariates for network inference and treated as a separate group in the regression penalization (Table S1). The r^2^ value assigned to each regression model is used as a confidence score to indicate the goodness of fit of the model. In the vehicle and treatment networks, all interactions with an absolute β-value < 0.01 or an r^2^ value < 0.1 and the connections to the surrogate variables were excluded.

**Figure 1:**
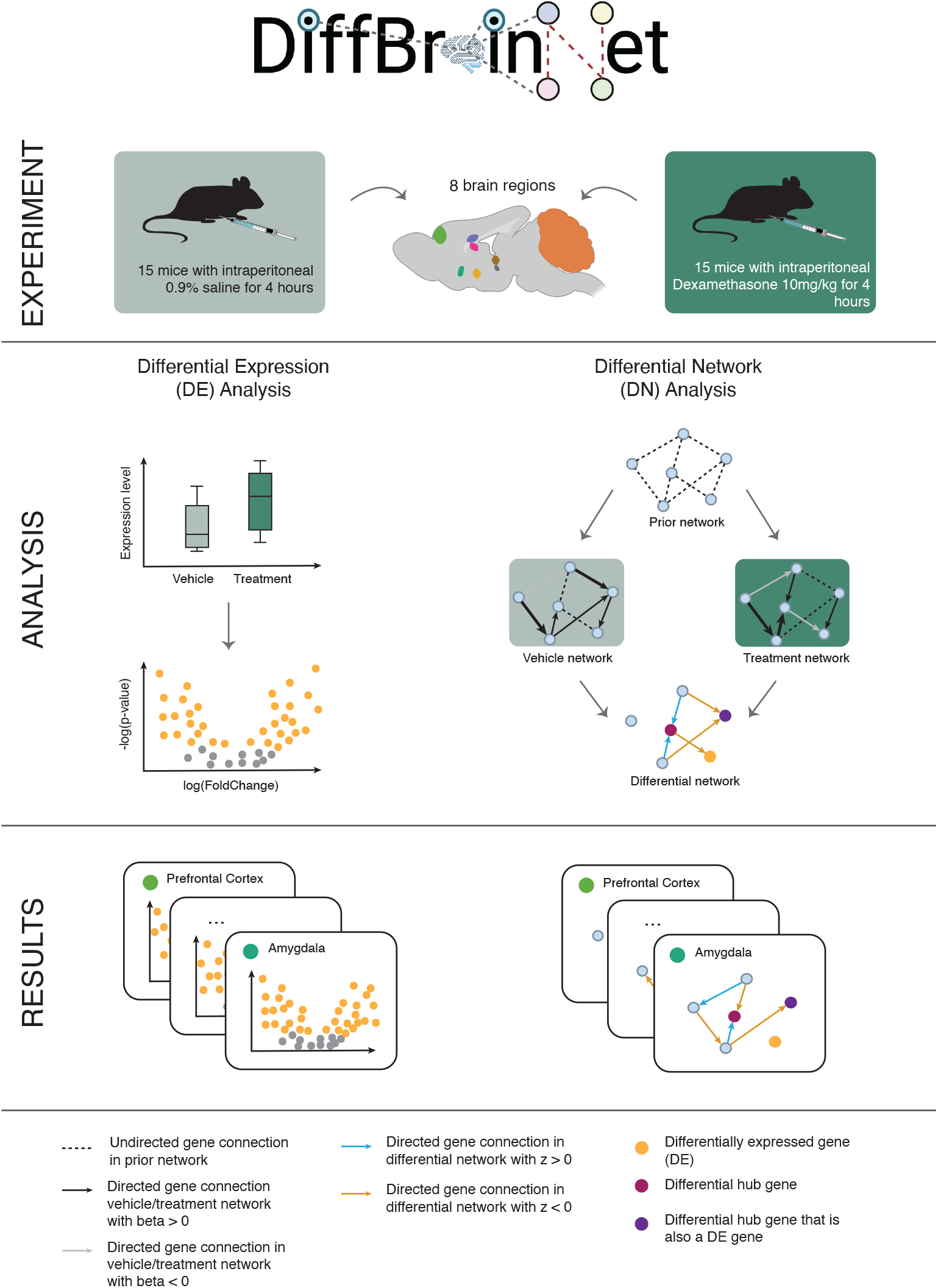
Schematic representation of experimental and analytical steps. DiffBrainNet is a resource of differential expression and differential networks in 8 mouse brain regions. (Experiment) *C57Bl/6* mice were treated intraperitoneally with 10mg/kg Dexamethasone or 0.9% saline as vehicle for 4hours. Eight different brain regions were isolated: amygdala – AMY, cerebellar cortex – CER, prefrontal cortex – PFC, paraventricular nucleus of the hypothalamus – PVN, dorsal *Cornu Ammonis* 1 – dCA1, ventral *Cornu Ammonis* 1 – vCA1, dorsal dentate gyrus – dDG, ventral dentate gyrus – vDG. (Analysis) We performed RNA sequencing in the 8 brain regions, followed by differential expression analysis (DE) and differential prior-knowledge-based genome-wide network analysis (DN). (Results) DiffBrainNet includes differential expression results and network results for all brain regions. DiffBrainNet logo was created with BioRender.com.

As a prior network we used FunCoup 5 [37], a database which contains about 6.7 million interactions between 19,771 genes in the mouse organism and that is provided as a framework to infer genome-wide functional couplings based on data of 10 different evidence types: physical protein interactions, mRNA co-expression, protein co-expression (based on the human protein atlas), genetic interaction profile similarities, shared regulation by transcription factor binding, shared regulation by miRNA targeting, subcellular colocalization, domain interactions, phylogenetic profile similarity, quantitative mass spectrometry data and gene regulatory data inferred from transcription factor bindings. FunCoup provides the edges of the basic network layout and KiMONo computes the weights of these edges fitted from the expression of the transcripts in each brain region and treatment paradigm.

#### Differential network analysis

A differential network (DN) for each brain region was calculated by combining the vehicle and treatment network using the DiffGRN approach [25] which describes differential relationships between two genes. Thereby, differential gene interactions were calculated from the regression’s β-values and their standard errors using a z-test:

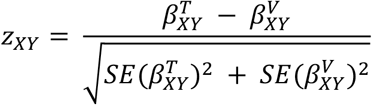

where β^T^_xy_ and β^V^_XY_ are the β-values of genes X and Y in the treatment and vehicle networks, respectively. A z-value > 0 indicates either a stronger positive correlation (0 < β^V^_XY_ < β^T^_XY_), a weaker negative correlation (β^V^_XY_ < β^T^_XY_ < 0) or a switch from negative to positive correlation (β^V^_XY_ < 0 < β^T^_XY_) between genes X and Y from vehicle to treatment network. A z-value < 0 indicates a stronger negative correlation (β^T^_XY_ < β^V^_XY_ < 0), a weaker positive correlation (0 < β^T^_XY_ < β^V^_XY_) or a switch from positive to negative correlation (β^T^_XY_ < 0 < β^V^_XY_) between genes X and Y from vehicle to treatment network. Z-values > 0 can be described as relative changes in gene expression leading to a more positive correlation (termed positive regulatory effect), while z-values < 0 can be described as relative changes in gene expression leading to a more negative correlation (termed negative regulatory effect) (S1B Fig). Differential interactions with an FDR adjusted p-value ≥ 0.01 associated with the z-score were excluded.

#### Hub gene analysis

We defined key regulators in the vehicle, treatment and differential networks, termed vehicle-, treatment-and differential-hub genes accordingly. The measure that we used to identify these key genes was the node-betweenness implemented in the igraph package, which describes the number of shortest paths going through a node [38]. Since we build the networks on top of a prior network, the node-betweenness in the networks (vehicle, treatment, differential) is driven by the prior network. We therefore normalized the node-betweenness as follows,

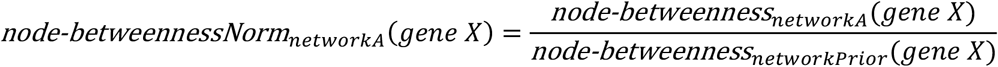

where node-betweenness_networkA_(gene X) is the node-betweenness of gene X in network A (e.g. DN of one brain region) and node-betweenness_networkPrior_(gene X) is the node-betweenness of the same gene X in the prior network. We defined all genes with a node-betweenness greater than 10,000 and a normalized node-betweenness greater than 1.0 as hub genes and compared them between brain regions as well as with the DE genes identified in the DE analysis.

### Gene set enrichment analysis

Enrichment of DE genes or differential hub genes was performed using FUMA GENE2FUNC [39] analysis based on Gene Ontology (GO, [40,41]), KEGG [42–44], Reactome [45] and genes carrying single nucleotide polymorphisms (SNPs) with genome-wide association to a variety of traits (analysis references the NHGRI-EBI GWAS Catalog [46] (https://www.ebi.ac.uk/gwas/) most recently updated on 18 September 2021). Default parameters were used in FUMA, with all genes expressed above threshold in all brain regions (n=12,830 genes) as the background list. To account for differentially sized input gene lists, only terms with at least 10% (unless stated otherwise) of the input genes overlapping with the term genes were considered and p-values were corrected using the Benjamini-Hochberg (FDR) method [47] to account for multiple comparisons. We used an FDR cut off of 5% for statistical significance.

### Shiny app

To make these data and analyzes searchable by all interested scientists, we created DiffBrainNet, which is accessible online at http://diffbrainnet.psych.mpg.de. The app was written in R (v4.0.5) [34], uses the shiny package (v1.7.1) [48] and several additional freely available packages (org.Mm.eg.db v3.14.0, shinythemes v1.2.0, ggplot2 v3.3.5, plotly v4.10.0, visNetwork v2.1.0, data.table v1.14.2, dplyr v1.0.7, stringr 1.4.0) and is hosted with ShinyProxy[49]. The source code of the app is available via github https://github.molgen.mpg.de/mpip/DiffBrainNet. The app can also be run locally using a docker image available on Docker Hub https://hub.docker.com/r/ngerst/diffbrainnet.

## Results

### DiffBrainNet: a brain-region specific resource and analysis framework for transcriptomic responses to glucocorticoid receptor activation

In this work, we set out to provide a resource of brain-region-specific transcriptome analyses at the gene-and network-level exploring the effects of a 4-hour 10mg/kg dexamethasone administration in 8 different mouse brain regions (Fig 1 top and S1A Fig). We used RNA sequencing to measure gene expression across the whole transcriptome and detected 12,976 genes across the 8 brain regions (exact numbers of transcripts per brain region in Table S1), with 12,830 genes being common across all 8 brain regions (Table S2).

Network analysis unravels the effects of relative gene expression changes that may not be detected at the individual DE genes. Therefore, gene expression networks for each condition per brain region were calculated with regression analysis based on a prior network using KiMONo [3]. As a prior network we used FunCoup 5.0 [37] which contains experimental data on about 6.7 million interactions between 19,771 mouse genes, of which 11,083 genes were also detectable in our dataset (5.4 million interactions). We inferred a DN per brain region by comparing the β-values of the regression analysis between the vehicle and treatment networks with a z-test, following the DiffGRN [25] approach. In addition, we also performed differential expression (DE) analysis to assess the gene-level responses to glucocorticoid receptor activation between vehicle and treatment (Fig 1 middle).

To examine if the DE genes are also the ones with the highest co-regulatory responses in the DNs we identified differential hub genes, i.e. genes with normalized node-betweenness above 1 (Fig 1 bottom). Furthermore, to identify pathways that are regulated by DE genes and/or differential hub genes we used enrichment analyses of GO terms, KEGG and Reactome pathways and GWAS significant genes. By applying this analysis framework, we were able to compare the transcriptomic responses across 8 brain regions on multiple complementary levels.

All data can be explored in an interactive online resource, called DiffBrainNet (http://diffbrainnet.psych.mpg.de). In the following, we illustrate results obtained from analyses using DiffBrainNet.

### Differential network analysis provides biological information beyond single gene-level analysis

We used our framework of combined DE and DN analysis to study the transcriptomic responses to glucocorticoids (GCs) across the eight brain regions in DiffBrainNet. Principal component (PC) analysis of the gene expression data showed that PC1 and PC2 explain 62% of the variance. The brain regions are separated by PC1 and PC2 whereas samples of the same brain region are comparable with respect to the first two PCs (Fig 2A). Treatment conditions were separated by PC4 and PC5 when PC analysis was applied on the samples of all brain regions together (Fig 2B). Over all 8 brain regions, we observed 2,092 DE genes (FDR adjusted p-value < 0.1) following dexamethasone administration of which 172 were shared DE across all brain regions (Fig 2C, Table S3). The majority of DE genes of each brain region were regulated in more than one region and only the minority (5.4-26.6%) was specific to a single brain region (S2A Fig, Tables S4-S11). The upregulated shared DE genes across all brain regions (N=129) were significantly enriched for biological processes related to cell death, response to stimulus, signal transduction and cell proliferation, whereas the downregulated ones (N=43) were enriched for developmental terms such as neurogenesis, cell differentiation and tissue morphogenesis (S2B Fig, Table S12).

**Figure 2:**
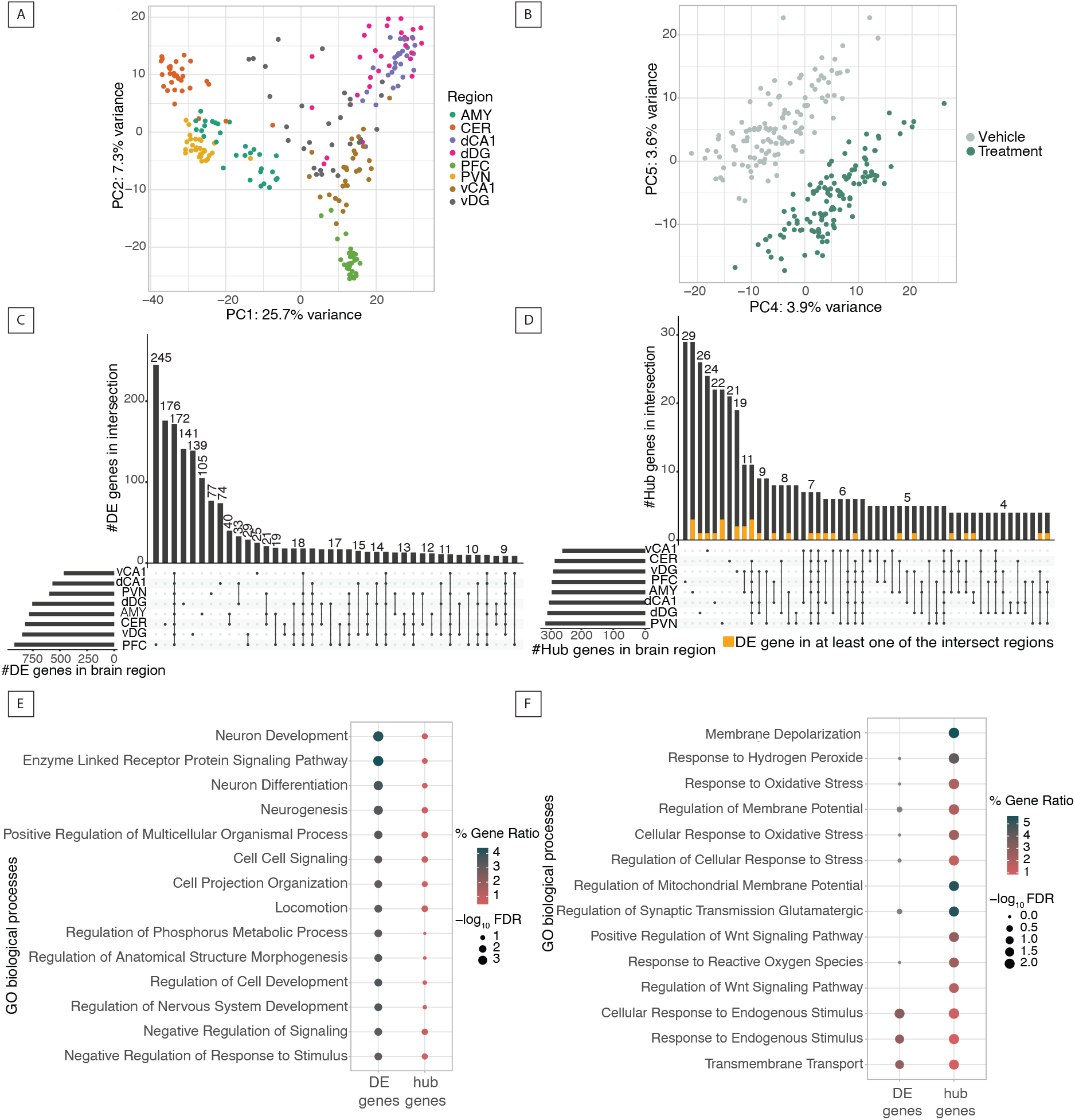
Differential network analysis provides distinct biological information from differential expression: the case of PFC. (A) Principal component (PC) analysis plot of PCs 1 and 2 explaining variance associated with brain region. (B) PC analysis plot of PCs 4 and 5 explaining variance associated with treatment group. (C) Upset plot comparing differentially expressed genes with FDR adjusted p-value smaller than 0.1 across 8 brain regions. (D) Upset plot comparing differential hub genes with a normalized node-betweenness above 1.0 across 8 brain regions. Proportions of intersection size bars coloured in yellow indicate genes that are also significantly DE genes in at least one of the intersection’s brain regions. (E) Dot plot for the top 14 GO terms most highly enriched for the unique DE genes and (F) for the unique differential hub genes in the PFC. (GO terms enrichment analyses are done with at least 10% of the input genes having to overlap with the genes of the term.)

In addition to DE analysis we performed DN analysis across the 8 brain regions and compared numbers and enrichment patterns of differential hub genes. We observed a total of 755 differential hub genes. The majority (over 73%) of these differential hub genes were shared between at least 2 brain regions (Fig 2D and S2C, Tables S13-S20), however, there were 7 differential hub genes shared across all investigated brain regions (*Sox5, Lpar1, Thy1, Mcam, Nell2, Rab3c, Zic1*) (Fig 2D and S2D, Table S21 and S22). Of all the 755 differential hub genes, only 174 were also DE genes in any brain region.

To further explore how DE genes and differential hub genes may relate to different biology, we compared the unique sets of these genes for the PFC, which was the brain region with the largest fraction of unique DE genes (n=920 total DE genes of which 245 (26.6%) were unique to PFC, Fig 2C, Table S11). PFC, together with AMY, was also the brain region with the highest fraction of unique differential hub genes (n=293 total differential hub genes of which 29 (9.9%) were unique in PFC, Fig 2D, Table S13). None of these 29 unique differential hub genes was also a DE gene in the PFC. A GO enrichment analysis on the unique DE and unique differential hub genes of the PFC respectively indicated that the biological functions related to these two sets of genes are distinct (Tables S23 and S24). While the biological processes with the highest enrichment for unique DE genes were mostly related to development and signaling (Fig 2E), the top terms for the unique differential hub genes were mainly global terms related to response to stress or stimulus (Fig 2F; n=14 terms). This suggests that DE and DN analyses reveal different but complementary information about the transcriptional response to the stimulus.

To show the added value of DN analysis we focused on *Abcd1*, a member of the ABC protein family known to actively transport GCs [50,51]. *Abcd1* is the unique differential hub gene that has by far the highest normalized node-betweenness in the PFC (normalized node-betweenness = 5,829, second highest is 4,013 for *Slc39a3*, Table S13) and many differential correlations, though it is not a PFC DE gene (FDR= 0.935; Fig 3A). However, in its DN there are 4 PFC DE genes (FDR < 0.1) and 7 genes that have a nominal DE p-value < 0.05 (Fig 3B, Table S25). By focusing at the pathways level, enrichment analyses of the DN of *Abcd1* supports a more general role of ABC transporters in the response to GCs (Fig 3C and D, Table S26). In addition, *Abcd1* is directly or indirectly connected to two other differential hub genes, *Tm7sf2* and *Pex5l*, suggesting that it is related to large interconnected DNs (Fig 3B). These smaller changes in the expression of genes that have in common their connectivity with *Abcd1* culminate in this gene’s status as a differential hub gene above the significance threshold, in spite of its too-subtle change at the individual expression level. Since biologically it is established that no gene works independently within a cell, these findings highlight the added value of network analysis to unravel distinct but complementary aspects of transcriptomic responses that can lead to specific molecular pathways identification.

**Figure 3:**
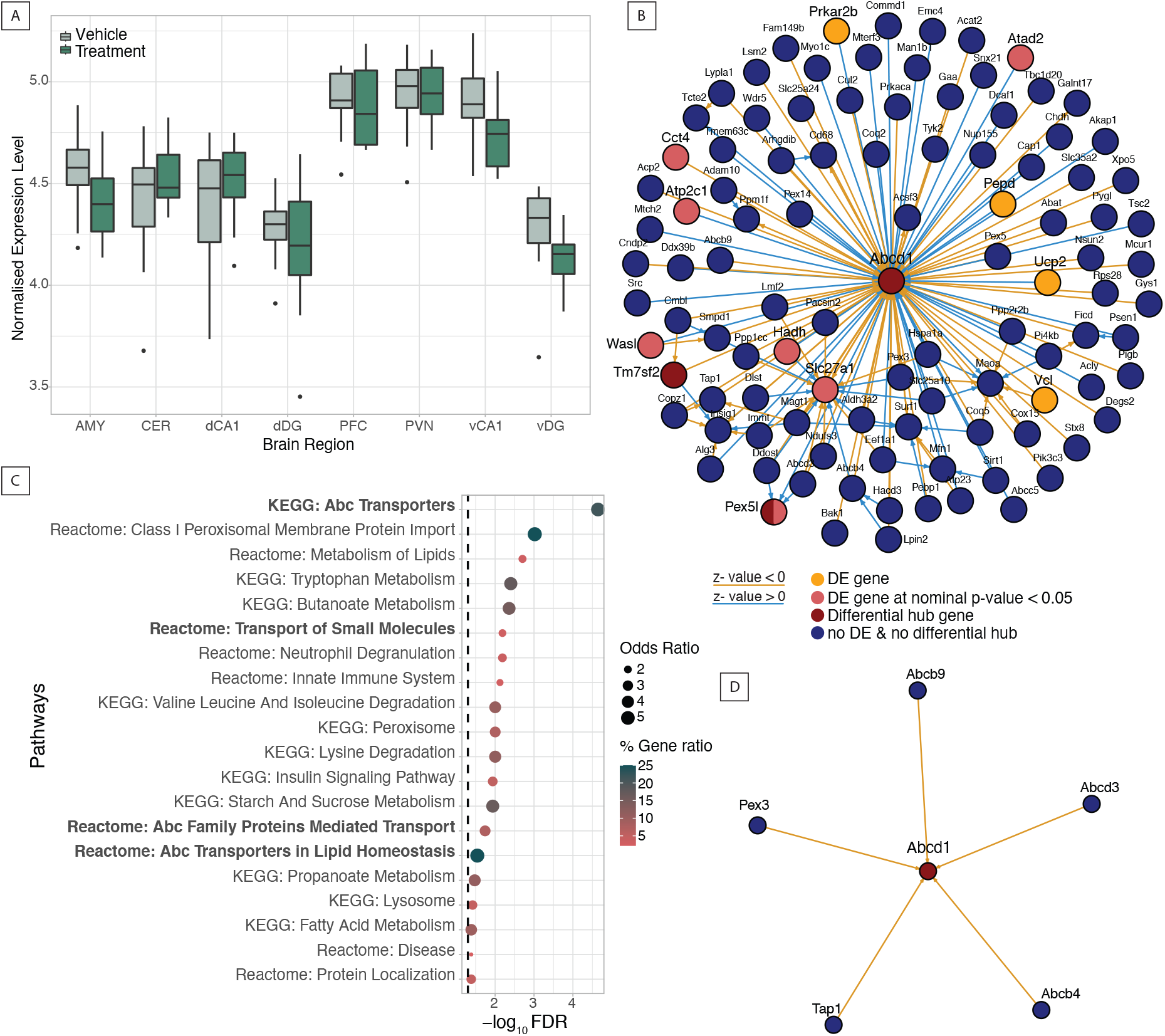
ABC transporters mediate dexamethasone response in the PFC at the network level. (A) Normalized expression of *Abcd1* in all brain regions at vehicle and after dexamethasone administration. *Abcd1* is not differentially expressed in any of the 8 regions. (B) *Abcd1* gene neighborhood in the differential network of the PFC. (C) KEGG and Reactome pathway enrichments for *Abcd1* and its differential neighbors in PFC. Bold labeled terms highlight a more general involvement of the ABC transporters pathway in the PFC response to glucocorticoids. (D) Network representation of the ABC transporters differential pathway.

### Differential network analysis supports the biological understanding of differentially expressed genes

Our next aim was to utilize DN to add an extra layer of interpretation to DE results, especially when the number of DE genes is insufficient for direct pathway analysis, indicating that the individual gene-level effects are very small. The vCA1 region of the hippocampus had the least number of unique DE genes from all brain regions with only 5.4% (n=25) of the total vCA1 DEGs (n=466) being unique to this region (Fig 2C and 4A and Table S8). Enrichment analysis at the GO level did not yield enriched terms (S3 Fig, FDR < 0.05 and Table S27). We next used these 25 unique DE genes as seeds in DiffBrainNet and found their differential neighbors, resulting in a DN containing 745 nodes, the 25 unique vCA1 DE genes and 720 differential neighbors (Table S28). This DN was enriched for genes associated via GWAS with autism spectrum disorder and depleted from genes associated with schizophrenia and general cognitive ability (Fig 4B and Table S29). These genes were now significantly enriched for GO terms associated with nervous system processes, cell morphogenesis, ion transport and synaptic signaling (Fig 4C and Table S30). This indicates that very small effects on multiple genes resulted in altered molecular connectivity in vCA1. This was not detected at the gene-level with DE analysis, but it was detected at the network-level with DN analysis.

**Figure 4:**
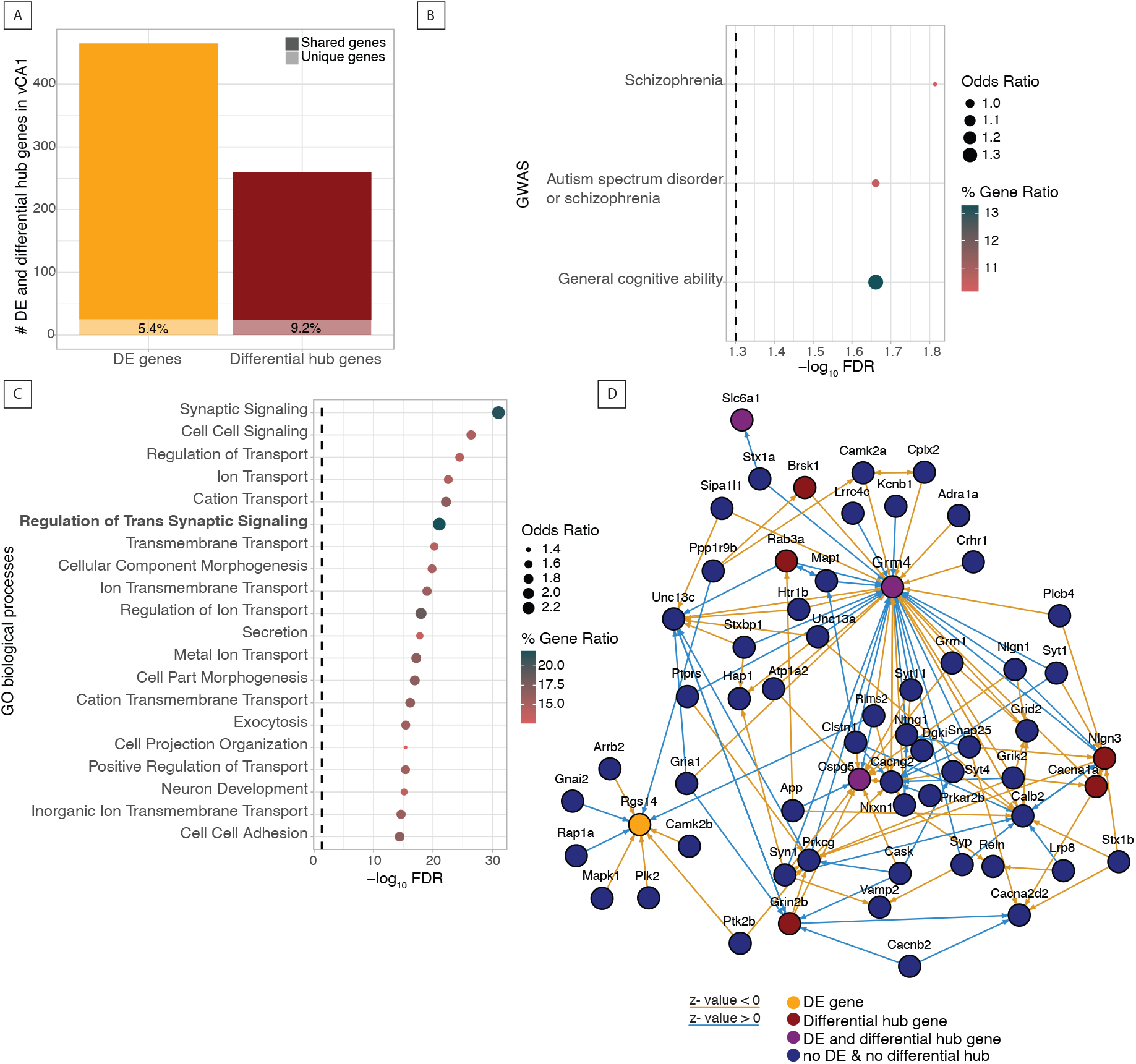
Differential network analysis supports the biological understanding of differential expression: the case of vCA1. (A) Number of unique and shared DE genes in vCA1 and number of unique and shared differential hub genes in vCA1. vCA1 has the least unique DE genes but the third highest percentage of unique differential hub genes of the eight brain regions. (B) Unique vCA1 DE genes and their differential neighbors are enriched for genes that carry SNPs associated with the GWAS traits schizophrenia, autism spectrum disorder or schizophrenia and general cognitive ability. (C) GO biological processes enrichment analysis of unique vCA1 DE genes and their neighbors. (D) Differential neighborhood of the genes that are part of the GO term regulation of trans-synaptic signaling and connected with the vCA1 unique DE genes. (GO terms enrichment analysis is done with at least 10% of the input genes having to overlap with the genes of the term. GWAS enrichment analysis is done with 16 of the input genes having to overlap with the genes of the term.)

We next focused on the top enriched GO term based on the gene ratio, “regulation of trans-synaptic signaling” (FDR = 8.71×10^−22^), and visualized the genes that were both part of the vCA1 DE genes network and associated with this term (Fig 4D). At the center of this DN was *Grm4*, which encodes a metabotropic glutamate receptor. *Grm4* showed many differential connections to other differential hub and DE genes including *Cacna1a*, which encodes a subunit of voltage-dependent calcium channels important for communication between neurons and synaptic signaling [52]. This trans-synaptic signaling network responded to dexamethasone by a number of changed correlations including several differential hub genes, beyond *Grm4* and *Cacna1a*, namely *Cspg5, Brsk1, Nlgn3, Rab3a* and *Grin2b*. This combination of DE and DN analysis was instrumental to identify potential biological responses to dexamethasone in the vCA1 region that were not readily detectable through DE analysis alone.

### DiffBrainNet can support exploring network changes related to candidate genes

We next sought to use our resource and analytical framework to investigate biological processes and pathways regulated by genes previously associated with risk for psychiatric disorders. DiffBrainNet provides the opportunity to study how genes of interest are co-regulated in different brain regions at vehicle-treated and after a stimulus, in this case glucocorticoid exposure. Here, we focused on understanding which biological processes were co-regulated by *Tcf4* (*Transcription factor 4*), a gene encoding a transcription factor with genome-wide significant associations to a number of different psychiatric disorders including schizophrenia, major depressive disorder and autism spectrum disorders [53] and for which mutations have been shown to cause neurodevelopmental disorders like for example Pitt-Hopkins syndrome [54].

We used DiffBrainNet to better understand this interaction by investigating the biological pathways co-regulated by *Tcf4* in the DNs reflecting changes associated with GR activation. *Tcf4* showed significant DE with dexamethasone in three of the brain regions, the amygdala, the vDG and the dDG, but in all brain regions the direction of change was the same (decrease following dexamethasone treatment; Fig 5A). While *Tcf4* did not show statistically significant DE in the PFC, previous work in this brain region using co-expression network analysis in human postmortem brain samples [55], has identified *Tcf4* as a master regulator in schizophrenia. When constructing a DN around *Tcf4* in the PFC, we identified 26 differentially connected genes including connections to DE genes (n=4) as well as differential hub genes (n=3, Fig 5B). The *Tcf4* PFC DN was enriched for genes that have been associated in GWAS with schizophrenia, autism and other neurobehavioral traits (Fig 5C and Table S31). This supports the observation that *Tcf4* networks are relevant for schizophrenia and adds the additional layer of the importance of *Tcf4* networks in the context of stress. Interestingly, the differential *Tcf4* network was not only enriched for GO terms related to development, but also autophagy and chromatin organization (Fig 5D and Table S32).

**Figure 5:**
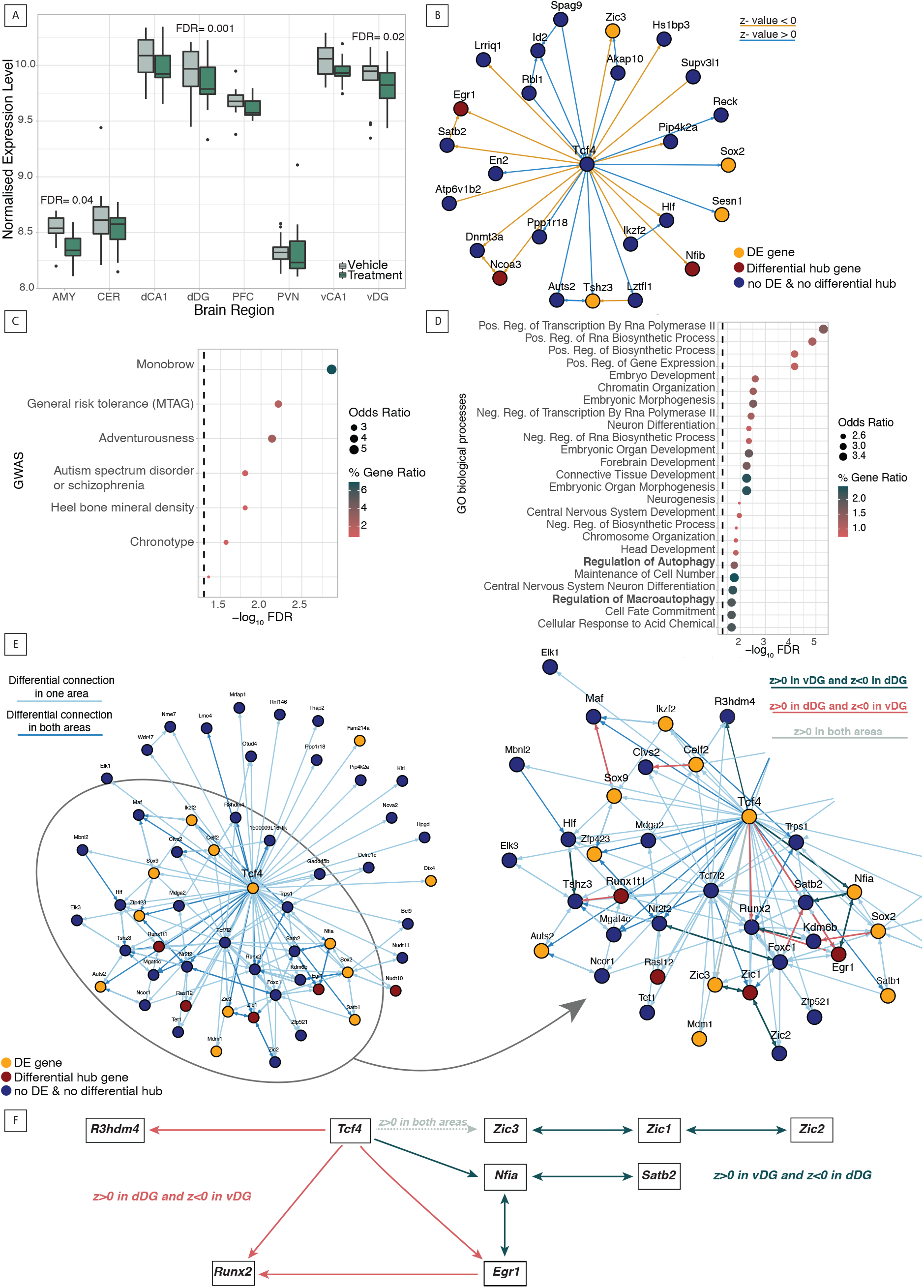
DiffBrainNet can support exploring network changes related to candidate genes: the case of *Tcf4*. (A) *Tcf4* is differentially expressed in the ventral and dorsal dentate gyrus (v/dDG) and in the AMY after 10mg/kg intraperitoneal dexamethasone treatment for 4 hours. (B) *Tcf4* DN in PFC. (C) *Tcf4* PFC DN is enriched for genes that carry SNPs associated with the GWAS traits schizophrenia, autism spectrum disorder or schizophrenia, adventureness and general risk tolerance among others. (D) GO biological processes enrichment analysis shows that members of the *Tcf4* PFC differential network are associated with development, neuronal differentiation, RNA biosynthetic processes and gene expression but also with regulation of autophagy (bold). (E) Differential network of *Tcf4* in both vDG and dDG (left). Zoom-in on a highly interconnected part of the DG *Tcf4* DN (right). Coloured with red are all the connections with a positive regulatory effect in dDG and a negative regulatory effect in vDG, coloured in black are all the connections with a negative regulatory effect in dDG and a positive in vDG and coloured in green is one of the connections that has a positive regulatory effect in both areas. (F) *Tcf4* molecular pathways that are co-regulated in an opposite manner in vDG and in dDG. *Tcf4* connections with the Zic transcripts and with *Satb2* and *Nfia* have a positive regulatory effect in vDG and a negative one in dDG whereas *Tcf4* connections with *Runx2, Egr1* and *R3hdm4* have a negative regulatory effect in vDG and a positive in dDG. (Enrichment analyses are done with at least 10% of the input genes having to overlap with the genes of the term.)

In contrast to the PFC, *Tcf4* was significantly downregulated in the dorsal and ventral dentate gyrus (Fig 5A). *Tcf4* is highly expressed in the hippocampal formation from the end of prenatal life and throughout adulthood [53]. We now aimed to use DiffBrainNet to investigate whether *Tcf4* being differentially expressed in the vDG and dDG of the hippocampal formation would have specific effects on each sub region’s molecular connectivity. From the 55 members of the *Tcf4* vDG and dDG DNs (Fig 5E), 20 are known *Tcf4* targets and/or protein interactors, according to the CHEA and TRANSFAC transcription factor targets databases [56,57] and the Pathway commons protein-protein interactions datasets [58]. An additional 11 genes are predicted *Tcf4* targets according to the MotifMap [59] and TRANSFAC [57] (S4 Fig and Table S33) (datasets assembled by the Harmonizome database, [60]). While most of the differential connections in this network were regulated in the same direction in both the vDG and the dDG, we also observed specific differential connections (n=24) that were regulated in an opposite manner between the two brain regions (Table S34 and selected ones in Fig 5F). *Tcf4* connections with the group of Zic genes, *Zic1, Zic2* and *Zic3*, suggested a positive regulatory effect (see Methods for explanation of term and S1B Fig) in vDG and a negative regulatory effect in dDG. Zic genes have been reported to play an important role in body pattern formation via the Wnt pathway [61], a pathway that has been extensively associated with *Tcf4* [62,63]. In addition, *Tcf4* had a positive regulatory connection with *Runx2*, another Wnt pathway effector [64], in dDG and a negative regulatory connection with it in vDG, suggesting that dexamethasone may mediate *Tcf4* effects on the Wnt pathway in a DG sub region-specific way. These types of analyses represent a thorough approach to hypothesis generation for further follow-up experiments of these effects.

## Discussion

The information provided by transcriptomic studies is far richer than a list of differentially expressed genes. Here, we have derived RNA expression from 8 mouse brain regions at vehicle and treatment (GCs) conditions and present DiffBrainNet, a resource and analytical framework, that provides access to DE and DN results. DiffBrainNet allows for direct synthesis and comparisons of the transcriptional landscape of all 8 brain regions at all conditions (Fig 6A). DiffBrainNet permits the search of DE genes unique to one brain region or common to any region combination at multiple FDR and fold-change cutoffs, the generation of plots and the chance to download the data (Fig 6C and 6D). In addition, DiffBrainNet offers the possibility to visualize the control (vehicle-treated), treatment (dexamethasone-treated) and differential networks in a single brain region and in any region combination, the ability to compare hub genes on all treatment levels at multiple node-betweenness thresholds and to download the network plots and data (Fig 6B and 6D).

**Figure 6:**
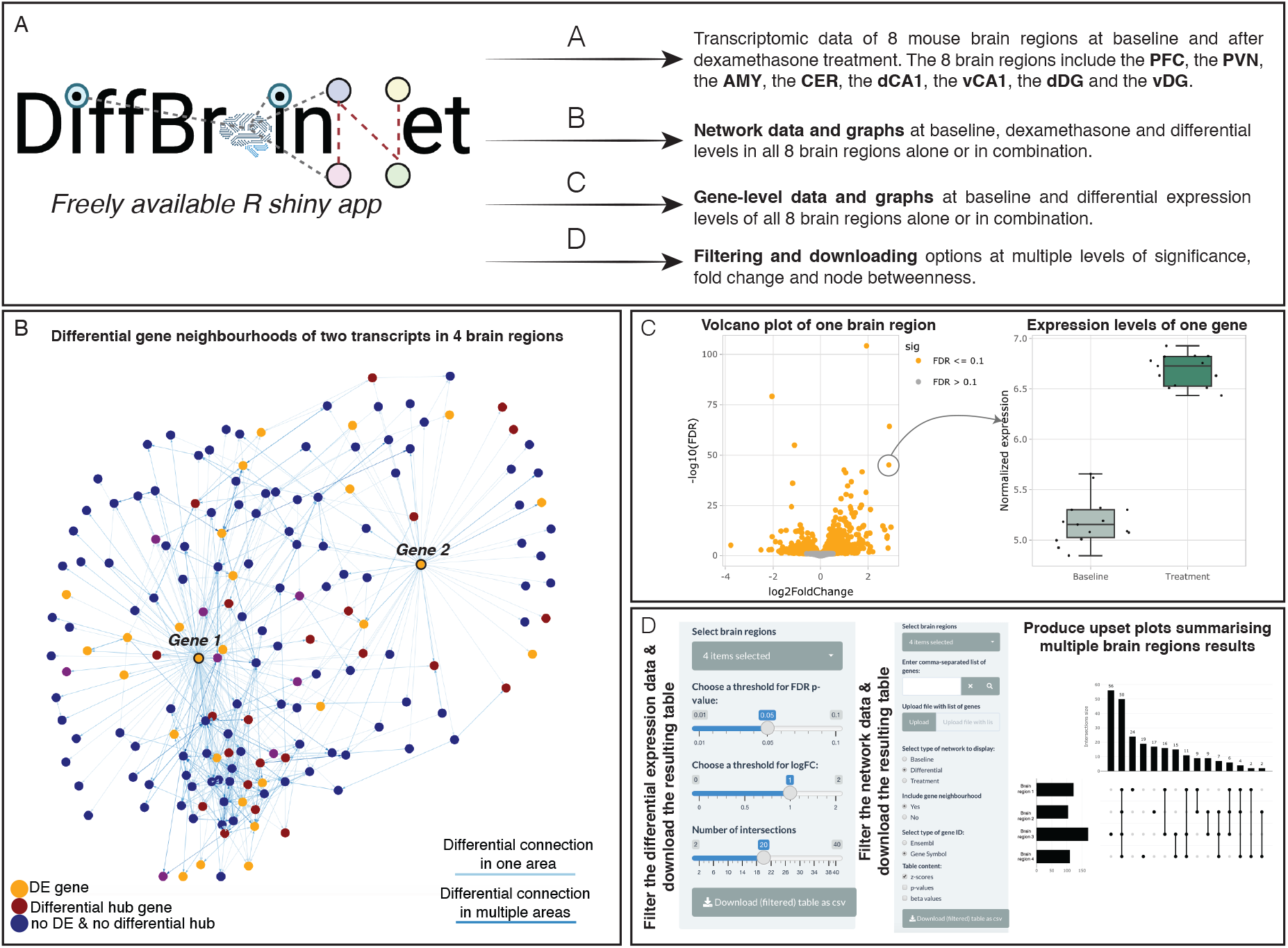
DiffBrainNet: a resource of gene expression and network data for 8 mouse brain regions. (A) DiffBrainNet includes gene expression and network data for 8 mouse brain regions at baseline, dexamethasone and differential levels. (B) DiffBrainNet provides network data for all 8 brain regions alone or in combination at baseline, treatment and differential levels. The data can be downloaded and plotted in the app. (C) DiffBrainNet provides gene expression data for all 8 brain regions. The data can be downloaded and plotted in the app. (D) The data both at the network-and the gene-levels can be downloaded using different thresholds of significance, fold change and node betweenness.

Comparing networks between two conditions is associated with a number of issues, as highlighted by De la Fuente [4] and described below. Comparison of networks uses mainly the node degree, which is a measure of a gene’s number of connections in two networks. This approach is highly dependent on the threshold that is used for the edges that are included in the two different networks and has proven challenging, since it is unclear how to choose comparable thresholds for two different networks. We sought to overcome this challenge by computing a single DN. We established a two-step method in order to differentially analyze networks. First, we used KiMONo to compute prior knowledge-based networks at vehicle-treated and following dexamethasone administration (treatment) conditions. Second, DNs were computed using DiffGRN. DiffGRN uses a z-test to calculate differential gene interactions based on the regression β-values of gene pairs at the vehicle and treatment network (S1B Fig) [25]. This approach provides differential interactions, thus eliminating the problem of having to compare two networks. This way we could pinpoint not only which genes but also which interactions of specific genes mediate the network changes. Moreover, by using prior-knowledge guided networks (KiMONo) [3], in which the expression of each gene is modeled by using the genes/proteins connected to it in a prior network as possible predictors in the regression model, we could compute vehicle and treatment networks of the same topological layout. This allowed for an even more robust comparison and reliable calculation of the z-values for the DN.

The use of biological knowledge in the form of a prior network, upon which the vehicle and treatment networks are built, is a substantial difference of KiMONo as compared to other network approaches such as weighted correlation network analysis (WGCNA) [2], which are built using correlation matrices without the use of prior-biological knowledge. DiffBrainNet is limited by a restricted search space, since it can only model interactions present in the prior network we chose to use. In the present analysis and the DiffBrainNet resource, we used FunCoup 5 to build our prior network [37]. FunCoup infers functional associations of genes or proteins using various data types and sources, including transcription factor binding sites, cellular and subcellular colocalization and protein-protein interactions. The use of such functional associations on the gene or protein level inferred by a variety of experimental data as prior-knowledge for predicting networks reduces the risk of false positives since the search space is restricted to known interactions and adds functional protein-level information to the transcriptomic data. Since, we provide the source code of all analysis (https://github.molgen.mpg.de/mpip/DiffBrainNet), a suitable prior network according to each research question can be chosen thus providing flexibility and specificity in hypothesis testing. By using prior-knowledge, the network metrics (node-degree, node-betweenness, modularity) are influenced by the prior network. To overcome this, we used normalized node-betweenness for all our analyses, which is defined as the node-betweenness in the calculated network divided by the prior network node-betweenness.

The combination of both gene-and network level analysis enriches our understanding of transcriptomic data and of biological implications. We showed that differential prior knowledge-based network analysis can unravel different and complementary aspects of the transcriptomic responses to a treatment as compared to individual gene-level analysis (DE). For example, we showed that in the PFC neither of the differential hub genes were also DE genes and that DE and DN analyses revealed distinct aspects of the transcriptomic responses. The DE genes explained effects mainly on signaling and development whereas the members of the DN explained mainly the cellular responses to the stimulus, GCs which are the main stress hormones, and stress.

DNs can be used to resolve underlying biological responses that are not detected by DE analysis. We identified *Abcd1* as the top differential hub gene in the PFC, which was not detected as a DE gene itself. ABC proteins are actively transporting GCs, including dexamethasone across the blood brain barrier and the placenta [50,51]. ABC transporters, synaptic biology and neuropsychiatric phenotypes have been previously associated in the literature. *Abcd1*-deficient microglia have been correlated with synaptic loss and axonopathy [65] pointing to an *Abcd1*-dysregulated network association with synaptic signaling problems. *Abcb1*, another member of the ABC transporters family, has been associated with stress adaptation and potential mediation of stress-related psychiatric disorders phenotypes [66]. These findings highlighted that the exclusive analysis of transcriptomic data at the gene-level does not capture all aspects of the transcriptional response to a stimulus, and the DN analysis can unravel distinct but complementary aspects that can lead to specific molecular pathways identification.

Finally, networks can be used for hypothesis generation and testing by choosing a suitable prior network. This approach can be exploited to generate hypotheses regarding the interactive effects of environmental exposures and the molecular underpinnings of specific genes. Using DiffBrainNet we analyzed the effects of dexamethasone on the co-expression network of a major psychiatric risk gene, *Tcf4*, in 3 different brain regions. *Tcf4* is expressed in the cortex, the hippocampus and the hypothalamic and amygdaloid nuclei predominantly at the end of prenatal life decreasing to lower expression levels throughout adulthood [53] and was shown to regulate neural progenitor cell maintenance and proliferation [67]. Animal models of gain and loss of function of *Tcf4* have shown its relevance for cognition, sensorimotor gating and neuroplasticity [68]. In addition, gene x psychosocial stress interactions have been reported for *Tcf4* [69], but little is known about relevant molecular pathways and brain regions for this interaction. With DiffBrainNet we showed that *Tcf4* mediates GCs effects in two sub-regions of the hippocampal formation, ventral and dorsal DG, at the gene- and at the network-levels since it is DE in those but only at the network-level for the PFC where is not a DE gene. The PFC DN of *Tcf4* was enriched for terms that include autophagy. The connection of *Tcf4* and autophagy has been previously described in the literature [62] but this is to our knowledge, the first report of a potential role of *Tcf4* in stress-related regulation of autophagy. This approach can be extended to the investigation of a wide spectrum of different gene lists - produced by GWAS studies for example - both at vehicle and after glucocorticoid exposure in a brain region-specific manner using DiffBrainNet. The results can be used to design more focused experiments to resolve targeted molecular mechanisms implicated in the pathogenesis of brain disorders.

In summary, through DN analysis we were able to identify specific molecular connectivity patterns governing transcriptomic responses to glucocorticoids that are not unraveled when investigating the differential gene expression levels alone. In our dataset, we inferred DNs in 8 mouse brain regions including a detailed segmentation of the hippocampal formation. With this work, we introduce DiffBrainNet, a resource and an analytical framework that includes both gene expression data and prior-guided genome-wide networks in these 8 brain regions at control (vehicle-treated), following GCs stimulation and at the differential level. DiffBrainNet can be used to pinpoint molecular pathways important for the basic function and response to GCs in a brain-region specific manner. It can also support the identification and analysis of biological processes regulated by brain and psychiatric diseases risk genes at the control and differential levels. We made these complex datasets and analyses available to all interested researchers via DiffBrainNet (access: http://diffbrainnet.psych.mpg.de, Fig 6).

## Data availability

Raw and normalized gene expression data generated in this study are provided at GEO under GSE190712 (https://www.ncbi.nlm.nih.gov/geo/query/acc.cgi?acc=GSE190712). Differential expression and differential network data can be downloaded from our resource DiffBrainNet.

## Code availability

Data analysis scripts and scripts that were used to generate the manuscript figures is available via github: https://github.molgen.mpg.de/mpip/DiffBrainNet. The source code of the shiny app is available via github as well: https://github.molgen.mpg.de/mpip/DiffBrainNet_ShinyApp.

## Author contributions

ACK and NG are joint first authors, contributed equally to this work and are listed alphabetically. ACK designed and carried out experiments, performed enrichment analyses, provided critical intellectual input and generated the paper draft; NG performed the differential expression and network analysis, implemented the resource as R shiny app, provided critical intellectual input and generated the paper draft; CC designed experiments, provided critical intellectual input and revised the paper draft; SR pre-processed the sequencing data; BP deployed and hosted the shiny app; MVS designed and carried out experiments and provided critical intellectual input; SS performed libraries preparation; MRH supported project organization and experimental procedures; JKA and EBB contributed equally to this work and are joint corresponding authors. JKA and EBB conceived the idea, obtained funding, supervised the study, designed experiments and analysis pipelines, provided critical intellectual input and contributed to paper draft writing.

## Acknowledgments

We thank the Biomaterial processing and repository unit (BioPrep) of the Max Planck Institute of Psychiatry for their contribution on RNA isolation. We thank Jonas Hagenberg for his help on the implementation of the Shiny app. NG is supported by the Joachim Herz Foundation.

