## Supplemental Figures for "DiffBrainNet: differential analyses add new insights into the response to glucocorticoids at the level of genes, networks and brain regions"

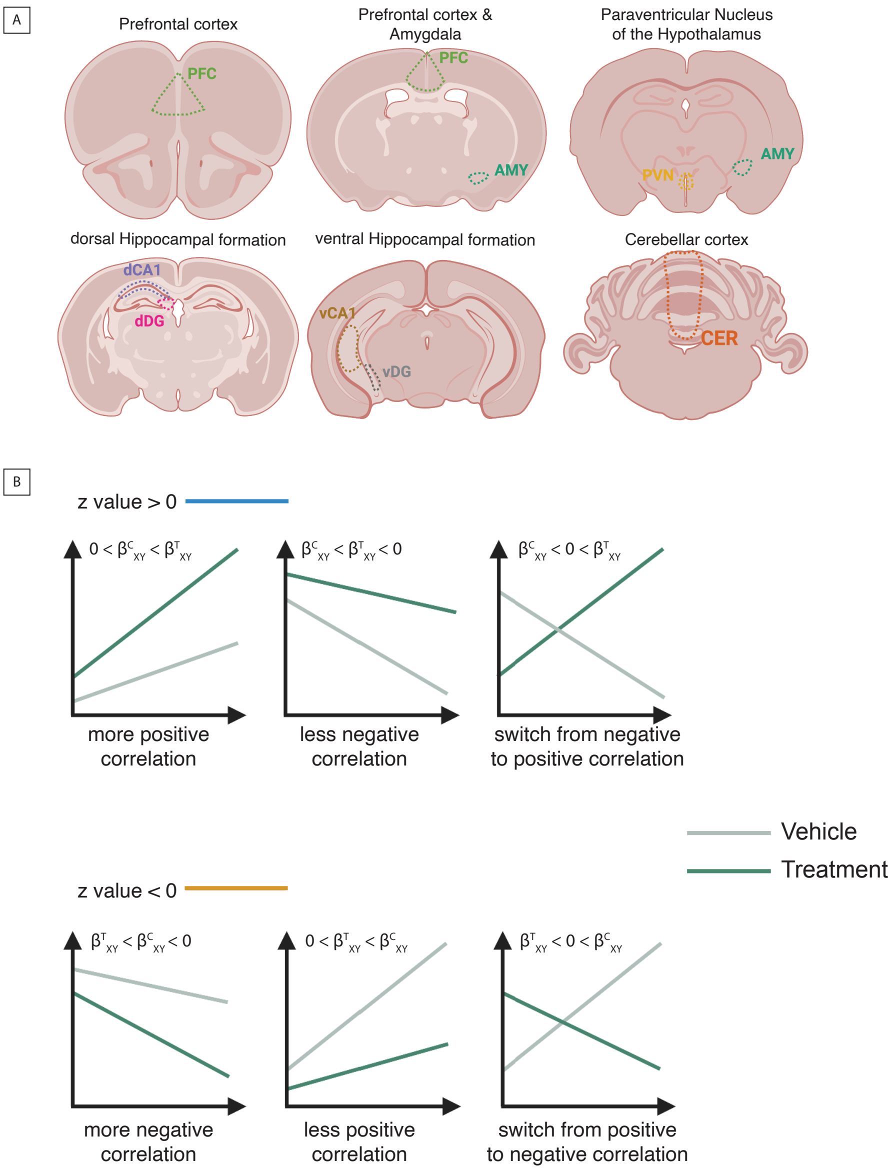


**Figure S1: Differential networks metrics explained.** (A) DiffBrainNet contains data for the prefrontal cortex (PFC), amygdala (AMY), paraventricular nucleus of the hypothalamus (PVN), dorsal and ventral *Cornu Ammonis* 1 (dCA1 and vCA1), dorsal and ventral dentate gyrus (dDG and vDG) and the cerebellar cortex (CER). (B) Z-values > 0 depict relative changes in gene expression leading to a more positive correlation, so an overall positive regulatory effect while z-values < 0 depict relative changes in gene expression leading to a more negative correlation, so an overall negative regulatory effect.

Mouse brain coronal sections were created with BioRender.com.

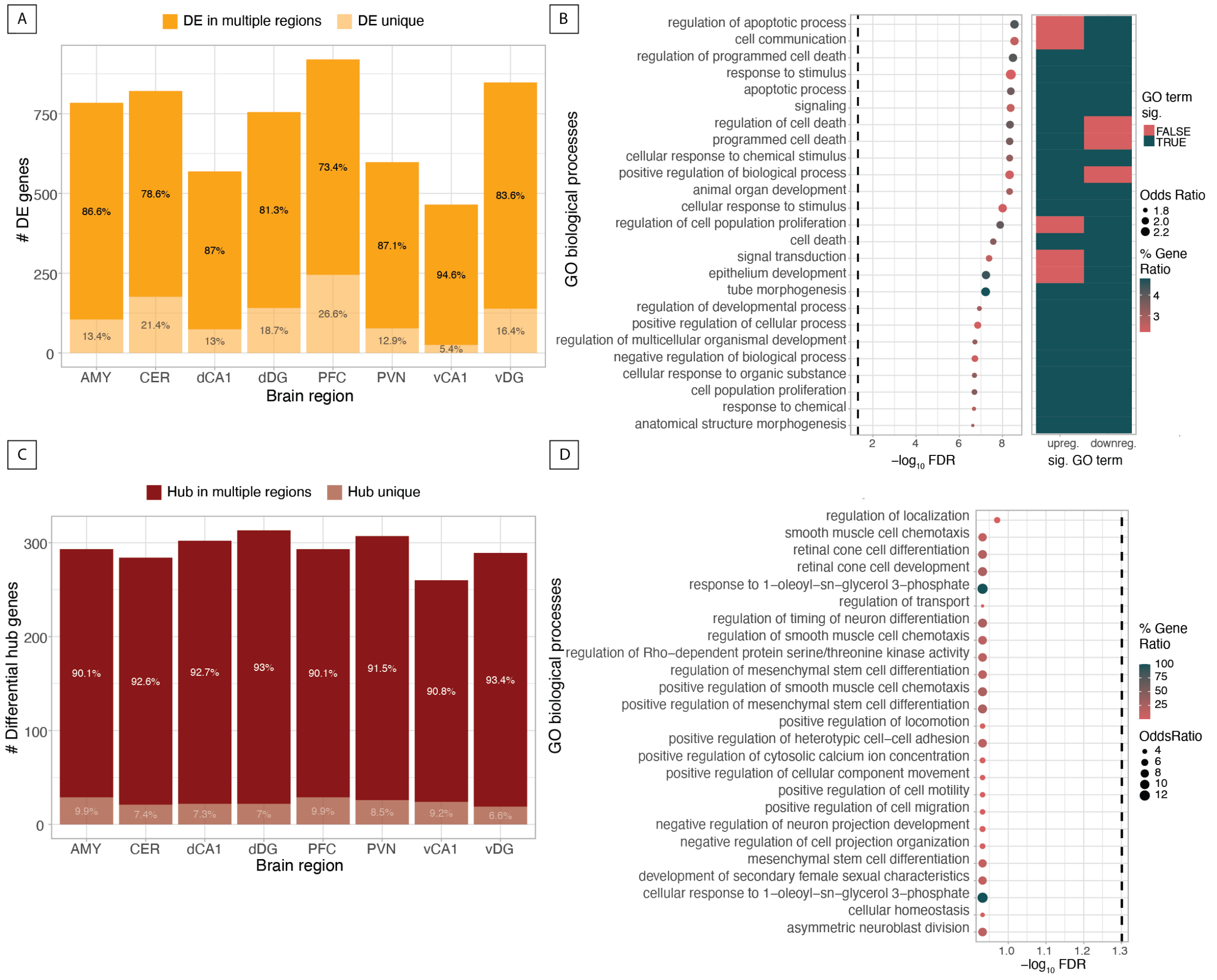


**Figure S2: Common response to dexamethasone of all brain regions.** (A) Percentage of unique and shared differentially expressed genes in each brain region. (B) Enrichment analysis of the 172 genes that are common differentially expressed genes in all eight brain regions. (C) Percentage of unique and shared differential hub genes in each brain region. (D) Enrichment analysis of the 7 genes that are common differential hub genes in all eight brain regions. Prefrontal cortex (PFC); paraventricular nucleus of the hypothalamus (PVN); amygdala (AMY); dorsal cornu Ammonis 1 (dCA1); ventral cornu Ammonis 1 (vCA1); dorsal dentate gyrus (dDG), ventral dentate gyrus (vDG) and cerebellar cortex (CER).

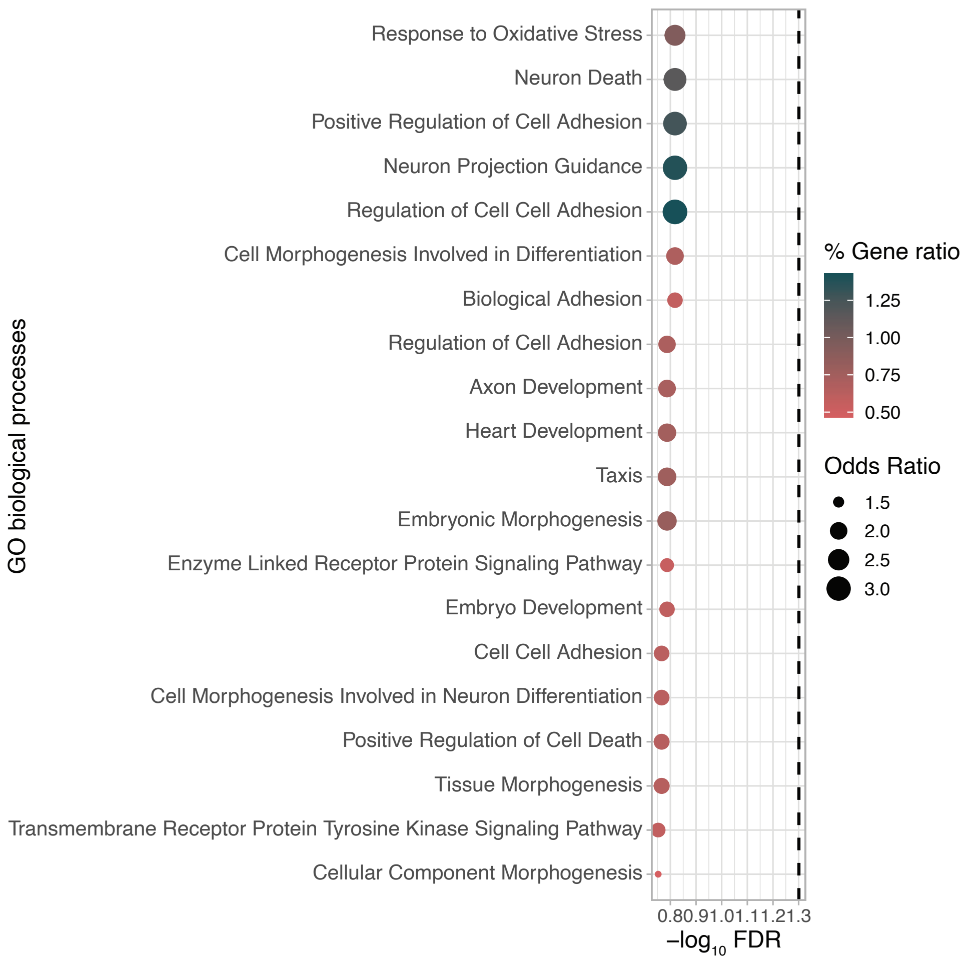


**Figure S3: vCA1 unique differentially expressed genes.** The vCA1 unique differentially expressed genes are not significantly enriched for any gene ontology terms. (GO terms enrichment analysis is done with at least 10% of the input genes having to overlap with the genes of the term.)

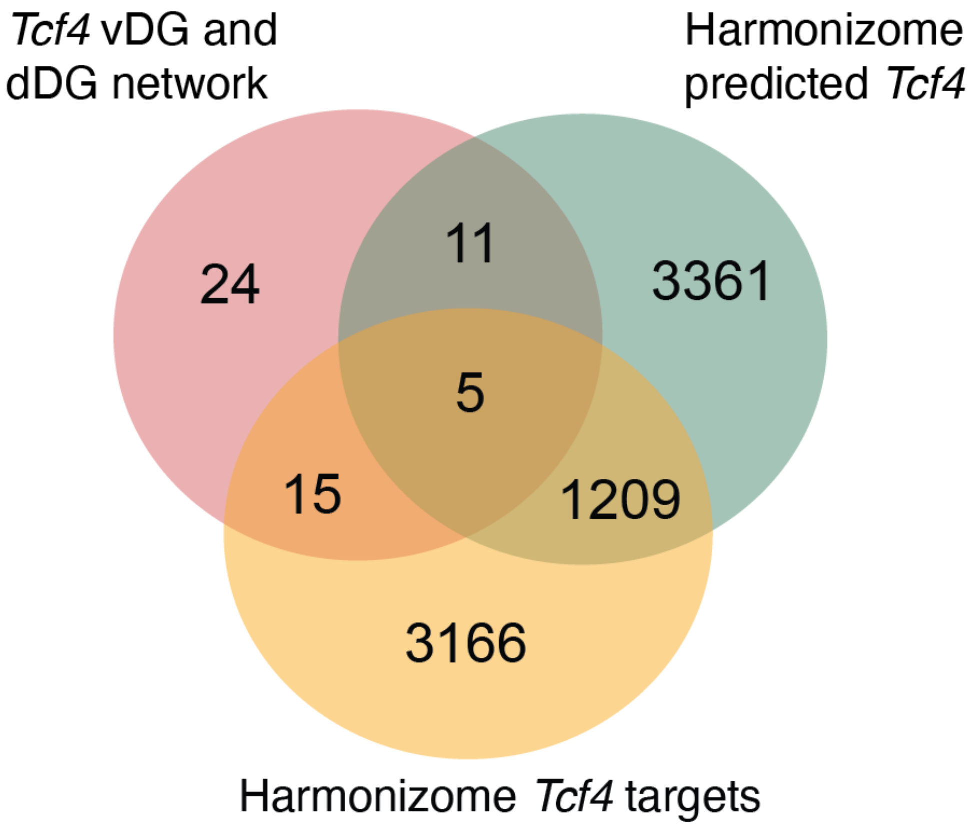


**Figure S4: *Tcf4* vDG and dDG network members.** Overlap of the genes that are members of the differential vDG and dDG *Tcf4* network with data from the CHEA, TRANSFAC and MotifMap transcription factor targets databases and from Pathway commons protein -protein interactions database collected from Harmonizome.
